# A possible genomic footprint of polygenic adaptation on population divergence in seed beetles?

**DOI:** 10.1101/2022.09.12.507575

**Authors:** Göran Arnqvist, Ahmed Sayadi

**Author notes:** Corresponding Author: Goran Arnqvist. “What we can measure is by definition uninteresting and what we are interested in is by definition unmeasurable” Richard Lewontin (1974).

## Abstract

Efforts to unravel the genomic basis of incipient speciation are hampered by a mismatch between our toolkit and our understanding of the ecology and genetics of adaptation. While the former is focused on detecting selective sweeps involving few independently acting or linked speciation genes, the latter states that divergence typically occurs in polygenic traits under stabilizing selection. Here, we ask whether a role of stabilizing selection on polygenic traits in population divergence may be unveiled by using a phenotypically informed integrative approach, based on genome-wide variation segregating in divergent populations. We compare three divergent populations of seed beetles (*Callosobruchus maculatus*) where previous work has demonstrated a prominent role for stabilizing selection on, and population divergence in, key life history traits that reflect rate-dependent metabolic processes. We derive and assess predictions regarding the expected pattern of covariation between genetic variation segregating within populations and genetic differentiation between populations. Population differentiation was considerable (mean F_ST_ = 0.23 - 0.26) and was primarily built by genes showing high selective constraints and an imbalance in inferred selection in different populations (positive Tajima’s D_NS_ in one and negative in one) and this set of genes was enriched with genes with a metabolic function. Repeatability of relative population differentiation was low at the level of individual genes but higher at the level of broad functional classes, again spotlighting metabolic genes. Absolute differentiation (d_XY_) showed a very different general pattern at this scale of divergence, more consistent with an important role for genetic drift. Although our exploration is consistent with stabilizing selection on polygenic metabolic phenotypes as an important engine of genome-wide relative population divergence and incipient speciation in our study system, we note that it is exceedingly difficult to firmly exclude other scenarios.

## Introduction

Some 160 years after Darwin (1859) suggested that natural selection plays a key role in the origin of new species, we now have a wealth of different types of evidence for Darwin’s general tenet (Schluter 2000; Coyne and Orr 2004). Yet, the relative importance of different types of phenotypic selection (e.g., natural vs. sexual selection) remains to be determined and we need to improve our general understanding of the nature of the molecular genetic basis of speciation and of the processes within populations that generate genomic divergence (Wolf et al. 2010). In this regard, one strategy is to study divergence between conspecific populations: studying selection in partially reproductively isolated forms can yield insights into selective processes before they become confounded by the accumulation of fixed differences between species (Via 2009; Burri 2017).

The large amounts of genome wide molecular genetic data generated by the last few decades of next generation sequencing efforts has made it possible to pursue “phenotype-free” bottom-up approaches in an effort to increase our understanding of incipient speciation. A common strategy in speciation genomics is to identify genes or regions of the genome that show unusually high differentiation between, for example, divergent populations. The physical location of detected outlier loci or their functional annotation is then used to make inferences about the processes responsible for divergence (Nosil and Feder 2012; Cruickshank and Hahn 2014). The more general supposition of speciation genomics is that incorporating analyses of genetic variation segregating within populations can provide information about the selective processes that generate divergence between populations (Wolf and Ellegren 2017).

Traditional methods based on outlier detection strategies can sometimes successfully detect the signature of selection if divergence is dominated by a limited set of major effect loci with independent effects and evolution proceeds by selective sweeps, whereby favorable variants are driven to fixation by divergent positive selection. However, issues are considerably more complicated when selection acts on ecologically relevant quantitative traits with a polygenic basis (i.e., complex traits). This is particularly true when net phenotypic selection (integrating over all life history stages, sexes, etc) is stabilizing and populations are adapting to different optima. Here, we expect populations to diverge through relatively minor shifts in allele frequencies across a very large number of loci showing epistasis for fitness (Gavrilets 2004; Höllinger et al. 2019) and such shifts are generally undetectable using outlier detection methods (Csilléry et al. 2018; Chevin 2019; Hayward and Sella 2021). In fact, because of redundancy, where a given optimal phenotype can be produced by an almost infinite number of allelic combinations across a large number of loci, even stabilizing selection which is uniform between populations is expected to generate suites of compensatory allelic changes across epistatically interacting loci. Such quasi-neutral polygenic drift is expected to result in population divergence, hybrid inferiority and, eventually, reproductive isolation (Gavrilets 2004; Fierst and Hansen 2010) (Figure 1).

**Figure 1.**
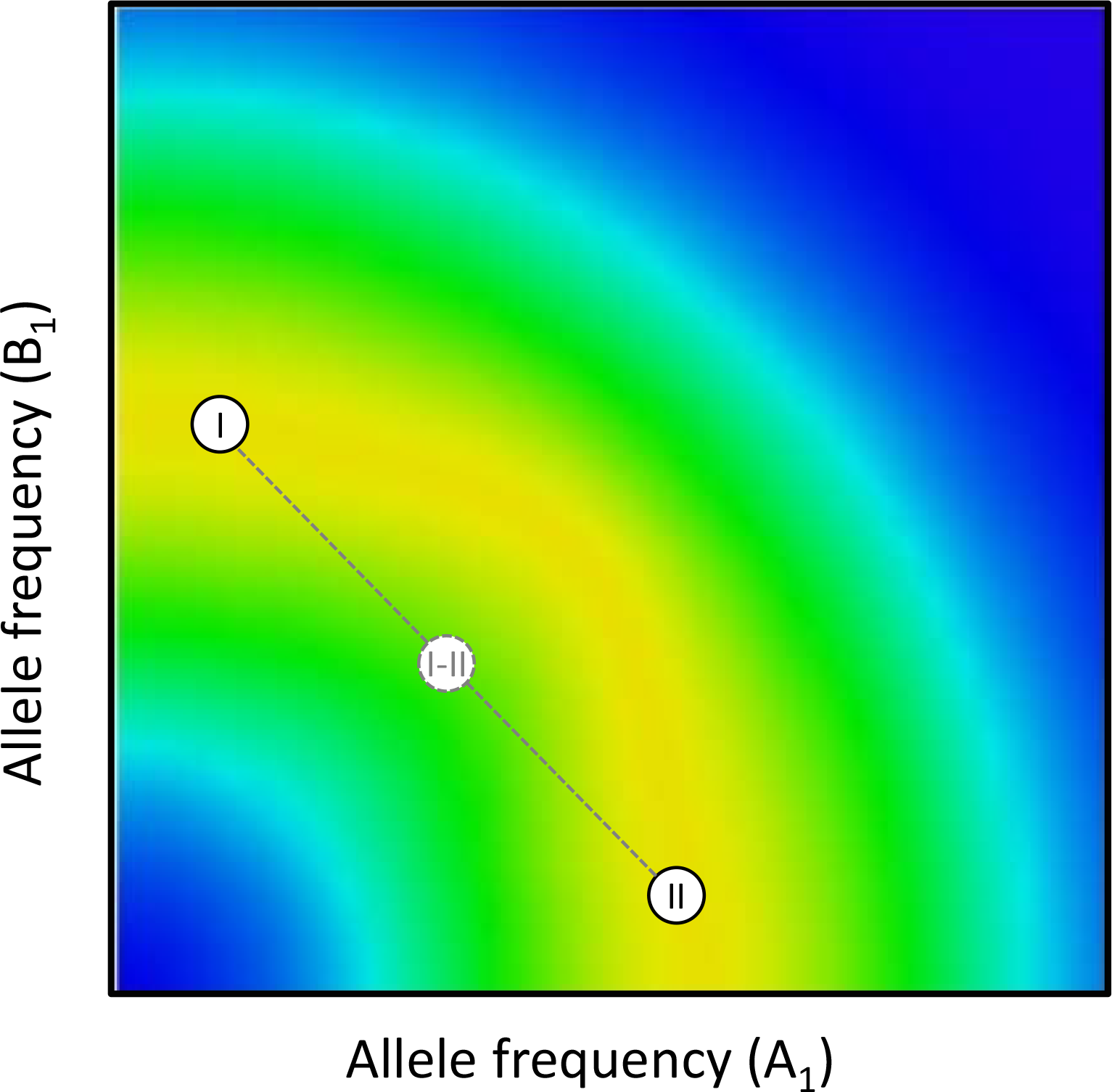
For polygenic traits under stabilizing selection, epistatic interactions between loci are inevitable (Barton 1986, 1989) because the fitness effect of an allele frequency shift in one locus will depend on the allelic state in many other loci (i.e., the genetic background). Because of complex multiplicative allelic effects on fitness across genes, fitness functions across loci tend to be curved (Rice 1998; Fierst and Hansen 2009). Here, there is a virtually infinite number of different combinations of allele frequencies across a large number of interacting loci that will all produce the same intermediate optimal phenotype. Populations may thus evolve along multidimensional “ridges” across holey adaptive landscapes (Gavrilets 2004). While population fitness may remain unaltered as populations move along such non-linear ridges, hybrid populations will tend fall in “holes” outside the maximum fitness ridges and show depressed fitness. This is illustrated in this simple 3D contour plot of allele frequencies in two loci where two populations (I and II) that occupy different locations along a curved maximum fitness ridge produce lower fitness hybrids (yellow denotes high and blue low fitness) (after Fierst and Hansen 2009).

Our empirical toolkit and theoretical understanding of speciation genomics is still to a large extent based on scenarios of selective sweeps involving one or a few independently acting or linked speciation genes (Nosil and Feder 2012; Cruickshank and Hahn 2014; Seehausen et al. 2014; Burri 2017; Wolf and Ellegren 2017). This state is unfortunate, given that this scenario is likely quite rare (Rockman 2012; Korte and Farlow 2013; Wellenreuther and Hansson 2016). We often expect (i) net selection on integrative quantitative traits such as life history traits to be stabilizing with intermittent shifts of the optimum and (ii) variation in such traits to have a highly polygenic basis, therefore inevitably showing marked epistasis for fitness (Fisher 1930; Barton 1986, 1989; Whitlock et al. 1995; Brodie 2000; Boyle et al. 2017; Barghi et al. 2020; Nosil et al. 2020). Moreover, the predicted role of selection in divergence critically depends upon genetic architecture (Chevin 2019). For example, loci under balancing selection are expected to contribute little to population divergence (Via 2009) and rather tend to show shared polymorphism and low divergence (Fijarczyk and Babik 2015) when assumed to act independently. In contrast, polygenic scenarios predict that a very large number of epistatically interacting loci encoding traits under stabilizing selection will, collectively, diverge either though selection and adaptation to shifting optima (e.g. Barton 1989; Höllinger et al. 2019) and/or through quasi-neutral polygenic drift (e.g. Gavrilets 2004; Fierst and Hansen 2010) (Figure 1).

Detecting the signal of polygenic adaptation in genomic data, however, remains a major challenge (Csilléry et al. 2018; Barghi et al. 2020; Hayward and Sella 2021). Here, our primary aim is to explore whether a role of stabilizing selection in polygenic population divergence can be unveiled by using an integrative and phenotypically informed approach, focused on analyses of genome-wide genetic variation segregating in divergent populations. We use a well-known insect model system where previous work has demonstrated a prominent role for stabilizing selection on key life history traits (e.g. Berger et al. 2016). We first consult multilocus theory of adaptation and divergence to ask what patterns we might expect if stabilizing selection on polygenic traits indeed plays an important role in population divergence in this system. We then assess whether these patterns are upheld in divergent populations. Importantly, a large body of previous experimental work in this system has generated insights into phenotypic selection and these insights allow us to form phenotypically informed predictions. We focus on broad scale genome-wide patterns of covariation across genes between standard indices of selection within populations (reflecting allele frequency spectra) and divergence between populations, rather than focusing on identifying single genes or regions by means of outlier detection strategies.

We base our analyses on population resequencing data from three well-studied populations of the seed beetle *Callosobruchus maculatus* (Bruchinae). These populations differ in geographic origin (see Materials and Methods) and are known to have diverged slightly but significantly in a series of life history phenotypes such as metabolic phenotypes (metabolic rate and respiratory quotient) (Immonen et al. 2016), body size (Rankin and Arnqvist 2008), juvenile growth rate (Arnqvist and Tuda 2010), egg-adult development time (Dowling et al. 2010), rate of reproductive senescence (Immonen et al 2016), ejaculate weight (Immonen et al. 2016) and composition (Goenaga et al. 2015), investment in immune function (Dougherty et al. 2017) and environment-specific reproductive fitness (Hotzy and Arnqvist 2009; Arnqvist and Tuda 2010). Reproductive incompatibility between populations is also significant but slight: egg-to-adult survival is depressed from some 90% within populations to approximately 80% in between-population crosses (Fricke and Arnqvist 2004; Rankin and Arnqvist 2008; Arnqvist and Tuda 2010). Previous studies show that net stabilizing selection on life history traits is pronounced in populations of this species (Berger et al. 2014, 2016). This derives in part from fundamental differences in male and female optima: briefly, males show a higher optimal pace-of-life than do females (Berger et al. 2014, 2016; Arnqvist et al. 2017). This is manifested as pronounced sexual dimorphism of rate dependent processes such as organismal metabolic rate (Berger et al. 2014; Arnqvist et al. 2017), accumulation of lipids during larval development (Lazarević et al. 2012), growth rate and development time (Arnqvist and Tuda 2010), adult activity (Berger et al. 2014) and adult life span (Fox et al.2003, 2004) and have resulted in persistent sexually antagonistic selection (Arnqvist and Tuda 2010; Berg and Maklakov 2012; Berger et al. 2014, 2016). Other studies have provided support for wide-spread balancing selection. For example, previous molecular genetic studies suggest an important role for balancing selection in maintaining polymorphism in life history genes in this species (Sayadi et al. 2019) and experimental work have identified sexually antagonistic pleiotropy as an important source of balancing selection (Berger et al. 2016; Grieshop and Arnqvist 2018). In line with this, Immonen et al. (2017) showed that genes with sex biased expression are enriched with those involved in metabolic processes and genes showing alternative splicing with sex biased expression of isoforms also show enrichment of metabolic genes.

### What might we expect?

Given the above, we derive eight predictions. (I) First, any polygenic trait under stabilizing selection will inevitably be affected by a large number of epistatically interacting loci. If, at one of these loci, a particular allele reaches high frequency in a given population then this will affect selection at other loci within the set (Figure 1). Shifts in allele frequencies across loci are thus predicted to be concerted, such that shifts in some loci will be associated with compensatory shifts in other loci (Hayward and Sella 2021), and divergence through this process is especially likely under weak epistasis (Fierst and Hansen 2010). The particular loci showing such shifts, whether caused by selection (Barton 1999), founder events (Arnqvist et al. 2014) or quasi-neutral polygenic drift (Fierst and Hansen 2010), are unlikely to be shared across diverging populations as stochastic events will have major effects on precisely which loci that show shifts. Reproductive isolation thus evolves because populations slowly evolve to new and unique combinations of allele frequencies (Barton 1989), resulting in the accumulation of Bateson–Dobzhansky–Muller (BDM) incompatibilities across many interacting loci (Gavrilets 2004; Fierst and Hansen 2010) (Figure 1). What pattern might this generate within and across diverging populations? Under a simple additive genetic architecture, long-term stabilizing selection on traits tends to erode underlying polygenic variation (Wright 1935). However, several plausible factors are capable of promoting polygenic variation, separately or in combination, by effectively generating balancing selection in loci encoding such traits. These include G × E interactions (Gillespie and Turelli 1989; Turelli and Barton 2004), antagonistic pleiotropy (Turelli and Barton 2004; Patten et al. 2010; Connallon and Clark 2014), overdominance (Turelli and Barton 2004), dominance reversal (Connallon and Chenoweth 2019; Grieshop and Arnqvist 2018), epistasis (Arnqvist et al. 2014) and frequency dependent selection (Mani et al. 1990; Turelli and Barton 2004; Bürger 2005). This is in accord with genetic studies suggesting that variants segregating at intermediate frequency contribute significantly to polygenic phenotypic variation (e.g. Charlesworth and Hughes 2000; Charlesworth et al. 2007; Mackay 2010; Charlesworth 2015; Chakraborty et al. 2018). Given the above, marked frequency shifts in a given locus, resulting in an excess of rare variants, is then expected to be compensated by relatively minor frequency shifts across many interacting loci. Importantly, because of redundancy, such shifts will very rarely be shared across populations. We suggest that, across the genome, genes that contribute to differentiation through this process should therefore generally tend to show signs of positive selection in one population (i.e., an excess of rare alleles) but belong to a class of genes showing signs of balancing selection in the other population (i.e., more even distributions of allele frequencies). We note that this expected apparent imbalance in selection contrasts with classic predictions for differentiation though divergent positive selection, where genes that build differentiation are expected to show signs of positive selection (for alternative alleles) in both populations.

(II) Previous work in seed beetles suggests that balancing selection is widespread among genes that are involved in metabolic pathways and thus dictate metabolic rate and sex specific pace-of-life phenotypes (see above). Since metabolic pathways involve products of many loci showing direct and indirect epistasis and because there should generally be stabilizing selection for intermediate fluxes, it has been predicted that balancing selection on metabolic genes should be common (Whitlock et al. 1995). We therefore predict that genes that show the predicted footprint of polygenic divergence, i.e. show an apparent imbalance in selection and simultaneously contribute to population differentiation, should be enriched with genes that are generally involved in metabolic pathways.

(III) More specifically, given that organismal metabolic rate is a key phenotype under stabilizing selection in our model system (see above), we predict that genes directly involved in regulating metabolic processes should show clear signs of the pattern described under the first prediction above.

(IV) Absolute population differentiation (i.e., d_XY_) is relatively insensitive to differentiation due to allele frequency shifts, as it ignores diversity within populations by essentially asking how different the most common alleles are in two populations. We therefore expect to see the pattern described under the first prediction above for relative (i.e., F_ST_) but not necessarily for absolute (i.e., d_XY_) divergence. Absolute divergence metrics are affected by a multitude of factors and processes and are difficult to interpret in isolation. Many scenarios based on divergent purifying selection predict that loci participating in population divergence should show both high relative and absolute divergence (e.g. Cruickshank and Hahn 2014). If balancing selection is prevalent, however, this will favor the maintenance of ancestral polymorphism in diverging populations while neutral ancestral polymorphisms are more likely to drift to fixation in diverging populations (Guerrero and Hahn 2017; Wang et al. 2019). The latter effect would tend to result in loci with less constraint showing relatively high absolute divergence.

(V) The considerations above suggest that the genetic basis for variation in any given polygenic life history trait should to some extent be distinct across populations, because of a substantial turnover in the genetic basis of traits during adaptation or quasi-neutral polygenic drift. Some loci should thus contribute more to trait variation in some populations and less in others, a point that reverberates the common observation of limited reproducibility in genotype-phenotype association studies of complex traits across populations of the same species (e.g. Chanock et al. 2007; Schielzeth et al. 2018). It has even been suggested that variation across loci and non-parallelism between replicated populations should be key hallmarks of polygenic adaptation (Barghi et al. 2020). We would thus predict that the degree of repeatability in relative population differentiation should be quite modest (Griffin et al. 2017), because differentiation should be built by partly different sets of loci across different sets of population pairs and the vector describing relative differentiation across genes should thus only partly be shared (Pritchard et al. 2010).Needless to say, however, this prediction is shared with other scenarios. Most importantly, divergence dominated by stochastic processes (e.g. genetic drift) can accommodate much the same pattern.

(VI) While we thus predict repeatability of relative population divergence to be modest at the gene level, due to redundancy, we predict that repeatability of population divergence in terms of functional classes or networks of genes should be considerable (Blount et al. 2018).

(VII) As stated above, for polygenic traits under stabilizing selection, drift or adaptation is expected to generate divergence though shifts in quasi-neutral allele frequencies across a large number of segregating loci (Barton 1999; Fierst and Hansen 2010). To further help distinguishing this scenario from divergence through drift across many independent neutral loci, one might predict that genome wide relative divergence is better explained by gene specific metrics that reflect the site frequency spectrum (e.g. Tajima’s D) than those that estimate selective neutrality by comparing segregating non-synonymous to synonymous variation (e.g., *π*N/*π*S) (Kryazhimskiy and Plotkin 2008).

(VIII) The probability that the frequency of a non-synonymous variant in a finite population will ultimately reach unity is a positive function of the strength of selection (Kimura 1964). For this reason, we often expect variants in large effect loci to show higher fixation probabilities. This dynamic is more complex for loci encoding polygenic traits under stabilizing selection that experience peak shifts. For example, Hayward and Sella (2021) showed that in many cases alleles with moderate effect sizes are instead predicted to show the greatest contributions to polygenic adaptation and that the fixation probability of such alleles is relatively high. This predicts that the relationship between fixation probability and effect size may be distinct in sets of epistatically interacting genes contributing to traits under stabilizing selection, compared with other sets of genes.

## Results

### General patterns of differentiation

The three populations, which were separated some 2000 years ago (see Materials and Methods), showed considerable overall relative differentiation (mean F_ST_, 95% bias corrected bootstrap CI; B-C: 0.248, 0.245 - 0.251; C-Y: 0.234, 0.230 - 0.236; B-Y: 0.256, 0.252 - 0.259), comparable in magnitude to previous estimates of differentiation between populations of seed beetles sampled in nature (Duan et al. 2016). To test prediction V, we asked to what extent gene specific population differentiation is shared between the three possible pair-wise comparisons (i.e., B-C, C-Y and B-Y). In order to allow explicit assessment of the repeatability of differentiation, we first performed a principal component analysis of the three vectors of pair-wise gene specific logit F_ST_ across all genes. This yielded a first PC which accounted for 49% of all variance in F_ST_ across genes and all three F_ST_ vectors loaded positively on this first PC (0.68, 0.71 and 0.71). A one-way ANOVA of all three values of gene specific F_ST_, using gene as a factor, yielded an intra-class correlation coefficient (repeatability) across the three comparisons of R = 0.25 (F_15305,30612_ = 1.94, P < 0.001). Gene identity thus explained some 25% of all the variance in F_ST_. The same exercise for log_10_ d_XY_ yielded a first PC that accounted for 89% of all variance in d_XY_ and all three d_XY_ vectors also loaded positively on this first latent variable (0.94, 0.95 and 0.95). The intra-class correlation coefficient here was R = 0.85 (F_19589,37752_ = 17.81, P < 0.001) showing that gene identity explained as much as 85% of all the variance in d_XY_. Moreover, the overall signals of relative and absolute differentiation were quite distinct, as the correlation across genes between the two vectors PC1_FST_ and PC1_dXY_ was as low as r = 0.15. We note that the correlation between F_ST_ and d_XY_ across genes was modest also for specific population pairs (B-C: r = 0.20; C-Y: r = 0.19; B-Y: r = 0.27).

These analyses are consistent with our prediction V, in that population pairs showed very restricted similarity in terms of which genes that exhibited high relative differentiation. Some ¼ of all variance in F_ST_ across genes was shared between the three comparisons, implying that ¾ is due to population specificity. For d_XY_, in contrast, more than ¾ of all variance was instead shared between the three pairwise comparisons. In an effort to understand if inherent properties of genes are associated with overall contribution to differentiation, PC1 was regressed against gene length and GC content. For PC1_FST_, this showed a positive association with both gene length (β’ = 0.15, *t* = 18.7, *P* < 0.001) and GC content (β’ = 0.08, *t* = 9.7, *P* < 0.001) but the model only accounted for 3% of the variance in PC1_FST_. For PC1_dXY_, neither the association with gene length (β’ = 0.013, *t* = 1.79, *P* = 0.073) nor with GC content (β’ = −0.011, *t* = −1.52, *P* = 0.127) was significant and the model only accounted for 0.03% of the variance. The addition of information of gene specific expression (i.e., FPKM values) did not significantly improve either of these two models (P > 0.09 in both cases). Hence, genes varied substantially and to some extent consistently in their contribution to absolute population differentiation, but gene length, GC content and gene expression were not strong predictors of their general role in differentiation.

We next asked if genes that generally showed highest overall absolute and relative divergence could be characterized functionally. Gene ontology enrichment analyses of the 400 genes with the highest values of PC1_FST_ showed significant enrichment for genes involved in a variety of biological processes, most of which involved various aspects of molecular transport and/or localization (SI Table 1). In particular, genes involved in transmembrane transport were overrepresented among the most divergent genes. The 400 genes with the highest values of PC1_dXY_, however, showed no apparent pattern of enrichment (SI Table 1). Moreover, PC1_dXY_ showed a positive relationship with *π*N/*π*S within the three populations (multiple regression; *F*_3,11459_ = 47.11, *P* < 0.001; all three *β'*s positive) while this relationship was strongly negative for PC1_FST_ (*F*_3,11266_ = 225.23, *P* < 0.001; all three *β'*s negative). This implies that genes generally showing high absolute differentiation are under less selective constraint while this pattern is reversed for genes showing high relative differentiation.

**Table 1.** The results of general linear models of the relationship between F_ST_ (logit transformed) between pairs of populations and segregating genetic variation within those populations, across all genes (model A1: *N* = 9735; A2: *N* = 8558; B1: *N* = 9888; B2: *N* = 8690; C1: *N* = 10150; C2: *N* = 8896). The value of (standardized) Tajima’s D_NS_ within populations alone explained a sizeable proportion of variance in F_ST_ across genes (model A1: 11%; B1: 13%; C1: 10%) and these effects remained strong when statistically controlling for GC content, gene length and the pN/pS ratios (model 2). Adding information on gene specific absolute expression (i.e., FPKM values) to model 2 did not significantly improve model fit (A2: *P* = 0.432; B2: *P* = 0.236; C2: *P* = 0.830) and neither did adding information on sex bias in gene expression (i.e., log2FC quartile) (A2: *F* _7,3607_ = 1.98, *P* = 0.053; B2: F7,3590 = 1.67, P = 0.111; C2: F7,3740 = 1.41, P = 0.195).

| A: $F_{ST}$ (Brazil - California) | | | | | | B: $F_{ST}$ (Brazil - Yemen) | | | | | | C: $F_{ST}$ (California - Yemen) | | | | | |
| --- | --- | --- | --- | --- | --- | --- | --- | --- | --- | --- | --- | --- | --- | --- | --- | --- | --- |
| Model | | $\beta$ | $SE_{\beta}$ | $t$ | $P$ | | $\beta$ | $SE_{\beta}$ | $t$ | $P$ | | $\beta$ | $SE_{\beta}$ | $t$ | $P$ | | |
| 1: | Brazil $D_{NS}$ | 0.057 | 0.023 | 2.52 | 0.012 | Brazil $D_{NS}$ | 0.230 | 0.021 | 10.71 | <0.001 | California $D_{NS}$ | 0.303 | 0.022 | 13.76 | <0.001 | | |
| | California $D_{NS}$ | 0.411 | 0.023 | 18.14 | <0.001 | Yemen $D_{NS}$ | 0.355 | 0.022 | 16.37 | <0.001 | Yemen $D_{NS}$ | 0.130 | 0.022 | 5.89 | <0.001 | | |
| | Brazil $D_{NS}^2$ | 0.060 | 0.018 | 3.38 | 0.001 | Brazil $D_{NS}^2$ | 0.008 | 0.017 | 0.47 | 0.640 | California $D_{NS}^2$ | -0.012 | 0.017 | -0.68 | 0.500 | | |
| | California $D_{NS}^2$ | -0.046 | 0.018 | -2.55 | 0.011 | Yemen $D_{NS}^2$ | -0.041 | 0.018 | -2.32 | 0.020 | Yemen $D_{NS}^2$ | 0.041 | 0.018 | 2.31 | 0.021 | | |
| | Brazil $D_{NS} \times$ California $D_{NS}$ | -0.550 | 0.021 | -26.07 | <0.001 | Brazil $D_{NS} \times$ Yemen $D_{NS}$ | -0.566 | 0.020 | -27.93 | <0.001 | California $D_{NS} \times$ Yemen $D_{NS}$ | -0.547 | 0.021 | -26.55 | <0.001 | | |
| 2: | Brazil $D_{NS}$ | 0.003 | 0.022 | 0.16 | 0.876 | Brazil $D_{NS}$ | 0.185 | 0.021 | 8.82 | <0.001 | California $D_{NS}$ | 0.273 | 0.021 | 12.80 | <0.001 | | |
| | California $D_{NS}$ | 0.368 | 0.022 | 16.42 | <0.001 | Yemen $D_{NS}$ | 0.318 | 0.021 | 14.79 | <0.001 | Yemen $D_{NS}$ | 0.084 | 0.022 | 3.89 | 0.022 | | |
| | Brazil $D_{NS}^2$ | 0.066 | 0.017 | 3.76 | <0.001 | Brazil $D_{NS}^2$ | 0.012 | 0.017 | 0.70 | 0.486 | California $D_{NS}^2$ | -0.013 | 0.017 | -0.77 | 0.444 | | |
| | California $D_{NS}^2$ | -0.032 | 0.018 | -1.79 | 0.074 | Yemen $D_{NS}^2$ | -0.041 | 0.017 | -2.34 | 0.019 | Yemen $D_{NS}^2$ | 0.056 | 0.017 | 3.20 | 0.001 | | |
| | Brazil $D_{NS} \times$ California $D_{NS}$ | -0.491 | 0.021 | -23.66 | <0.001 | Brazil $D_{NS} \times$ Yemen $D_{NS}$ | -0.527 | 0.020 | -26.39 | <0.001 | California $D_{NS} \times$ Yemen $D_{NS}$ | -0.480 | 0.020 | -23.98 | <0.001 | | |
|  | GC content | 0.002 | 0.004 | 0.48 | 0.630 | GC content | 0.007 | 0.003 | 2.12 | 0.034 | GC content | 0.002 | 0.003 | 0.51 | 0.608 |  |  |
|  | Gene length | 4.81E-05 | 4.60E-06 | 10.47 | <0.001 | Gene length | 3.99E-05 | 4.31E-06 | 9.25 | <0.001 | Gene length | 4.26E-05 | 4.38E-06 | 9.72 | <0.001 |  |  |
|  | Brazil pN/pS | -0.173 | 0.044 | -3.91 | <0.001 | Brazil pN/pS | -0.191 | 0.037 | -5.21 | <0.001 | California pN/pS | -0.451 | 0.047 | -9.65 | <0.001 |  |  |
|  | California pN/pS | -0.416 | 0.050 | -8.26 | <0.001 | Yemen pN/pS | -0.238 | 0.034 | -6.91 | <0.001 | Yemen pN/pS | -0.084 | 0.034 | -2.45 | 0.014 |  |  |

### Gene classes

On average, genes encoding male seminal fluid proteins showed a higher absolute differentiation relative to other genes, while this was not true for female reproductive proteins (Figure 2) (SI Table 3). Absolute differentiation was in fact significantly higher for male seminal fluid proteins then for female reproductive proteins (Welch’s *t*-test of log10 d_XY_; B vs. C: *t* = 3.05, *P* = 0.002; B vs. Y: *t* = 1.82, *P* = 0.069; C vs. Y: *t* = 2.62, *P* = 0.009). This difference was, however, not seen for relative population differentiation (Welch’s *t*-test of logit F_ST_; *t* < 1.40 and *P* > 0.16 in all three cases) (Figure 2).

**Figure 2.**
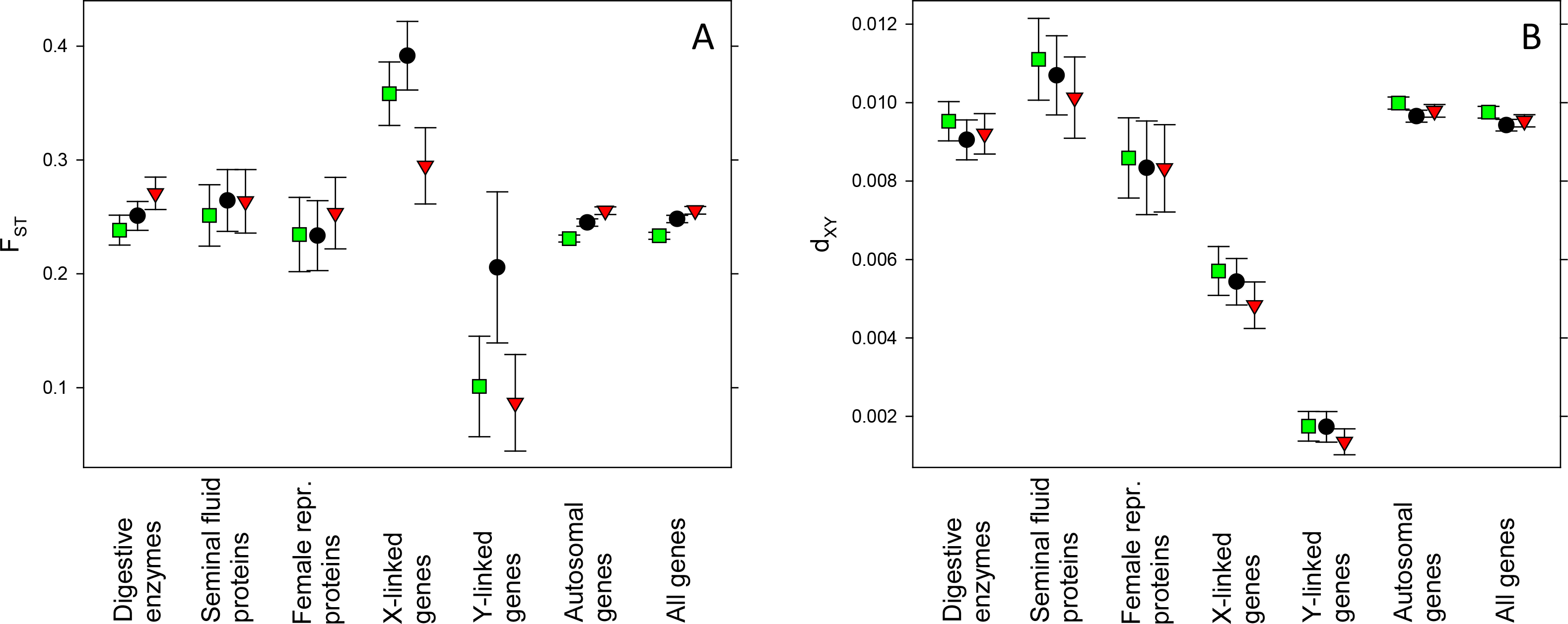
Overall population differentiation for functional sets of genes across three *C. maculatus* populations. Shown are mean (± bootstrap 95% CI) (A) relative and (B) absolute population differentiation for pairs of populations (green: C vs. Y; black: B vs. C; red: B vs. Y). (A) Relative divergence was highest for X-linked genes but was also somewhat elevated in digestive enzymes and male seminal fluid proteins. (B) Absolute divergence was markedly elevated in male seminal fluid proteins and was reduced in X- and Y-linked genes. See SI Table 3 for statistical evaluation.

Relative, but not absolute, mean differentiation was higher in genes encoding enzymes involved in digestion in larvae compared with other genes (SI Table 3) (Figure 2). A direct comparison with male seminal fluid proteins showed that these showed significantly higher absolute (Welch’s *t*-test of log10 d_XY_; B vs. C: *t* = 2.58, *P* = 0.009; B vs. Y: *t* = 1.32, *P* = 0.186; C vs. Y: *t* = 2.35, *P* = 0.019) but not relative (Welch’s *t*-test of logit F_ST_10; *t* < 0.35 and *P* > 0.72 in all three cases) differentiation compared with digestive enzymes.

Genes located on the sex chromosomes often, but not always, show higher rates of divergence compared with autosomal loci, which is seen as the combined result of their sex-biased inheritance, lower recombination rates and lower effective population size (Meisel and Connallon 2013; Wilson Sayres 2018). We found that mean relative differentiation was high among X-linked loci (SI Table 3, Figure 2). In contrast, absolute divergence was very low for X-linked loci and the elevated relative differentiation seen was primarily a result of the fact that X-linked genes in *C. maculatus* show low nucleotide diversity within populations (Sayadi et al. 2019). This contrast can thus serve to illustrate the fact that relative and absolute measures of population differentiation can sometimes show very different signals (Cruickshank and Hahn 2014). The low differentiation seen here for Y-linked genes is consistent with previous findings of relatively low nucleotide diversity and efficient purifying selection in most hemizygous genes located on the Y (Sayadi et al. 2019), which is presumably shared among populations to a large extent.

### Variation within and differentiation between populations

One of our goals was to test our predictions by examining global relationships across genes between the overall properties of site frequency spectra within populations to differentiation between populations. We did so in a series of explicit general linear models of the relationship between F_ST_ or d_XY_ between populations on one hand and metrics summarizing the gene specific pattern of segregating genetic variation within populations on the other. We were particularly interested in how common within population molecular estimates indices of selection, here represented by Tajima’s D (Tajima 1989) for non-synonymous sites and by *π*N/*π*S, related to estimates of population differentiation. Tajima’s D provides a classic and useful summary that relates the frequency spectrum of segregating sites to expectations under the neutral equilibrium model, where positive values correspond to an excess of common polymorphisms consistent with balancing selection and negative values are consistent with a recent selective sweep (Fijarczyk and Babik 2015). Its absolute value, however, is sensitive to demographic processes and significance testing of single estimates can be difficult. We note that our analyses are based on variation in Tajima’s D across genes, which all share a common recent demographic history within populations, and we do not engage in tests of significance of Tajima’s D.

The correlation between Tajima’s D for non-synonymous sites and log *π*N/*π*S across all genes was very low (B: r = −0.05; C: r = 0.01; Y = −0.04), illustrating that these two metrics summarize distinct aspects of the nature of segregating genetic variation within populations.

Our analyses uncovered several patterns. These models are summarized in Tables 1 and 2. Across all genes, Tajima’s D predicted both relative and absolute differentiation between populations, but did so in very distinct ways. Variation in F_ST_ was primarily determined by the interaction between Tajima’s D in the two populations, where genes showing positive Tajima’s D in one population but negative Tajima’s D in the other were those genes that overall contributed most to relative population differentiation (Figure 3; see SI Figure 1 for fitted function). This was true irrespective of whether variation in potentially confounding factors was controlled for or not (Table 1). In contrast, genes showing negative or positive Tajima’s D in both populations contributed the least to overall differentiation.

**Table 2.**
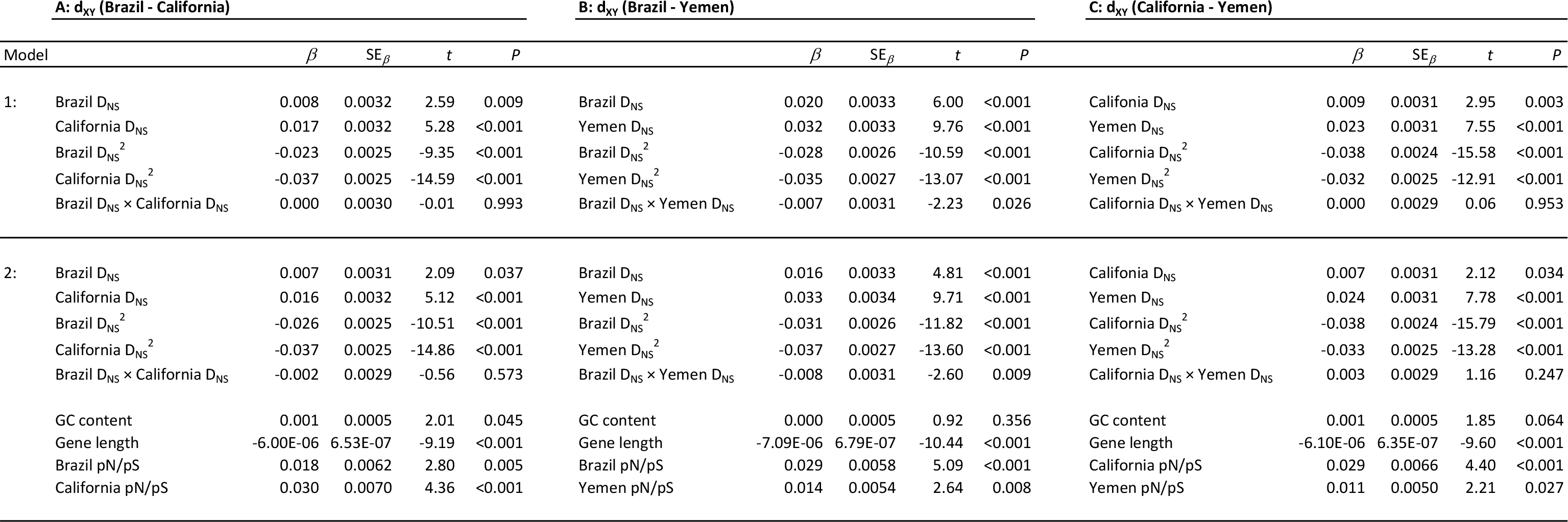
The results of general linear models of the relationship between d_XY_ (log10 transformed) between pairs of populations and segregating genetic variation within those populations, across all genes (model A1: *N* = 10096; A2: *N* = 8616; B1: *N* = 10289; B2: *N* = 8766; C1: *N* = 10547; C2: *N* = 8968). The value of (standardized) Tajima’s D_NS_ within populations alone explained a significant proportion of variance in d_XY_ across genes (model A1: 4%; B1: 5%; C1: 6%) and these effects remained when statistically controlling for GC content, gene length and the pN/pS ratios (model 2). Adding information on gene specific absolute expression (i.e., FPKM values) to model 2 did not significantly improve model fit (A2: *P* = 0.413; B2: *P* = 0.092; C2: *P* = 0.170). In contrast, the addition of information on sex bias in gene expression significantly impoved model fit in all three cases (i.e., log2FC quartile) (A2: *F* _7,3607_ = 10.2, *P* < 0.001; B2: F7,3590 = 9.79, P < 0.001; C2: F7,3741 = 8.72, P < 0.001).

**Figure 3.**
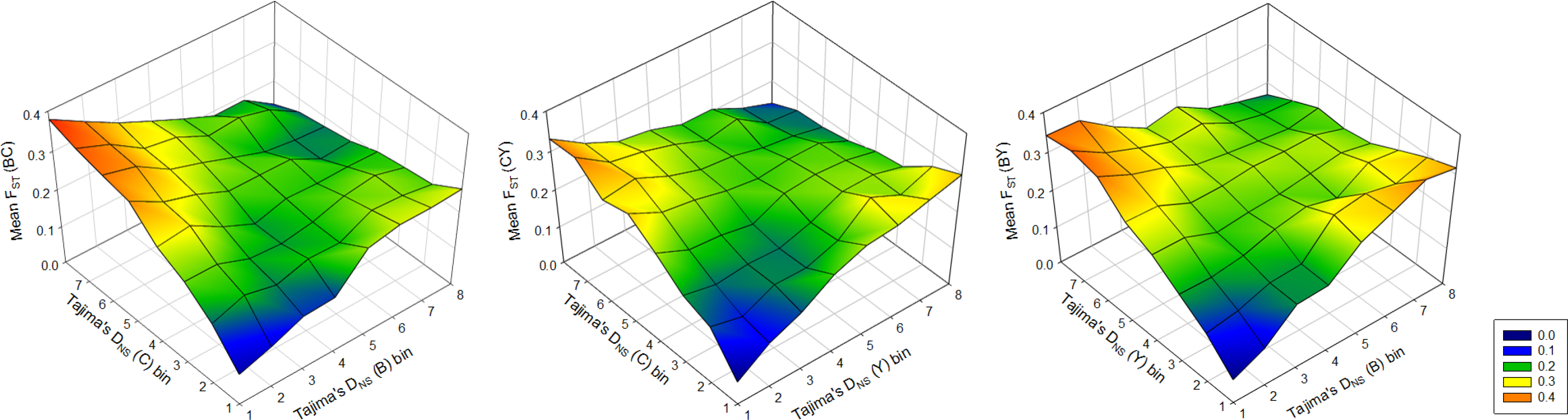
3D mesh plots of average F_ST_ across all genes as a function of Tajima’s D (based on non-synomymous SNPs) within populations. Here, genes have been binned into eight groups (N = 1293 – 1522 genes per bin; see SI Figure 4) by the value of Tajima’s D, to illustrate the fact that genes under positive selection in both populations (i.e., the most negative values of Tajima’s D) have the lowest values of F_ST_ overall. Genes showing an apparent signal of balancing selection in one population and positive selection in the other have the highest overall F_ST_ (see Table 1 for statistical evaluation). The comparison B-C is shown to the left, C-Y in the centre, and B-Y to the right.

The analyses of variation in Tajima’s D across genes are thus consistent with prediction I in showing that genes that exhibit an excess of rare variants in one population and a scarcity of rare variants in the other are those that contribute most to relative population differentiation (F_ST_), such that that an apparent imbalance in inferred gene specific selection between two populations was associated with differentiation between them. Next, we tested prediction II by asking whether such genes tend to be enriched with genes with particular functions. We selected those genes that showed an absolute value of the difference in Tajima’s D_NS_ > 2.5 in two populations (imbalance in inferred selection) while simultaneously showing F_ST_ > 0.3 between them (high relative differentiation). These three subsets of genes (B vs. C: N = 238; B vs. Y: N = 275; C vs. Y: N = 164) were then subjected to gene ontology enrichment analyses. Analyses of these sets (SI Table 1) revealed significant enrichment for a series of gene ontology terms, with a striking overrepresentation of genes involved in various metabolic processes (SI Figure 2). For example, significant terms occurring in at least two out of the three population comparisons (and with enrichment based on at least two genes in the set) included the parental biological process term “single-organism metabolic process” but also the children terms “ncRNA metabolic process” and “organonitrogen compound catabolic process”. Collectively, these analyses are thus consistent with prediction II and suggest that genes that show signs of balancing selection in one population but positive selection in the other and that simultaneously contribute to relative population differentiation tended to be genes involved in a variety of metabolic processes, including catabolism and biosynthesis (SI Figure 2).

In agreement with prediction VI, the above analyses show that although the repeatability of relative population differentiation was low at the gene level (see above) it was high at the level of broad functional classes of genes: 85% (28/33) of all significantly enriched terms involved metabolic processes (i.e., contained the term “metabolic”, “catabolic” or “biosynthetic”) (SI Figure 2).

To assess prediction III, which is founded in the fact that metabolic rate phenotypes have been identified as a key polygenic phenotype in this system, we first selected all genes annotated as being involved in processes that regulates the rate of metabolic organismal pathways (i.e., Biological Process GO:0019222) (in total N = 269). Mean logit F_ST_ was higher in metabolic rate genes compared with all other genes in all three population comparisons, and significantly so in two out of three cases (Welch’s *t*-test; B vs. C: *t* = 1.13, *P* = 0.26; B vs. Y: *t* = 2.70, *P* = 0.007; C vs. Y: *t* = 2.17, *P* = 0.031). More importantly, we replicated our three main models (i.e., model 1 in Table 1) using only this subset of genes, predicting that imbalances in Tajima’s D between populations would predict relative differentiation between them. These analyses showed that the interaction term between Tajima’s D indeed still had the strongest effect on relative differentiation between them (SI Figure 3; SI Table 2).Compared with models including all genes (Table 1), models of this subset of genes showed somewhat accentuated effects of the interaction term relative to all other effects in the model in two out of three comparisons (mean effect size [*f*^2^] for the interaction term, B vs. C: from 0.066 to 0.034; B vs. Y: from 0.074 to 0.088; C vs. Y: from 0.067 to 0.082) (Cohen 2013). Although not entirely conclusive, these analyses are at least broadly consistent with a predicted important role for metabolic rate genes in polygenic differentiation in this system.

The relationship between absolute differentiation between population pairs and indices of selection was very distinct from that of relative differentiation described above. Variance in d_XY_ was primarily determined by the quadratic terms of Tajima’s D in both populations (Table 2), such that genes showing intermediate and near-zero values of Tajima’s D in both populations were those genes that overall contributed most to absolute population differentiation (Figure 4). Models of variance in d_XY_ were less predictive of differentiation compared with those of F_ST_ but the pattern, again, was robust against the potential impact of confounding factors (Table 2). Moreover, the association between of *π*N/*π*S and d_XY_ described the inverse of that between of *π*N/*π*S and F_ST_ (see signs of coefficients in model 2 in Tables 1 and 2). Genes showing relatively high *π*N/*π*S ratios in both populations, consistent with lower selective constraint (Kryazhimskiy and Plotkin 2008), were those that contributed most to overall absolute differentiation. These analyses thus suggest that genes that exhibit segregating variation more consistent with the neutral expectation in both populations were those that contributed most to absolute population differentiation. This is consistent with a relatively more important role for genetic drift in generating absolute differentiation.

**Figure 4.**
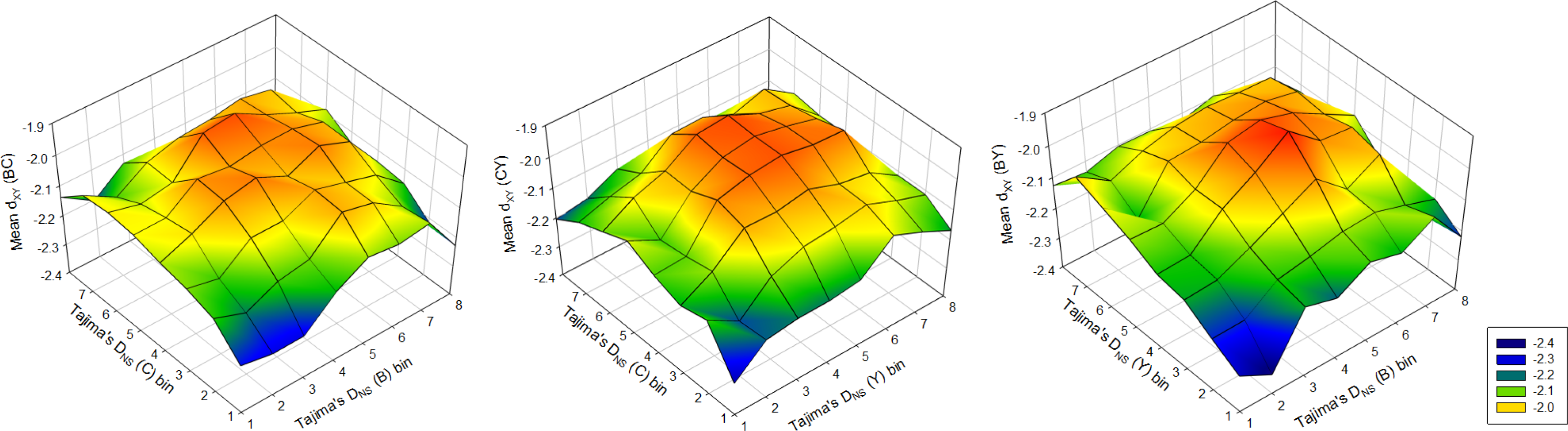
3D mesh plots of average dXY (log10 transformed) across all genes as a function of Tajima’s D (based on non-synomymous SNPs) within populations. Here, genes have been binned into eight groups (N = 1293 – 1522 genes per bin; see SI Figure 4) by the value of Tajima’s D, to illustrate the fact that genes showing the the greatest average absolute diverence are those with intermediate values of Tajima’s D in both populations. These are generally close to zero (see Table 2 for statistical evaluation). The comparison B–C is shown to the left, C–Y in the centre, and B–Y to the right.

We note that our models showed that gene-specific relative differentiation is better predicted by Tajima’s D than by *π*N/*π*S (Table 1), consistent with prediction VII. Gene length was associated with both relative and absolute differentiation, but was so in contrasting ways (Tables 1 and 2). While longer genes showed higher relative differentiation on average, they showed lower absolute differentiation.

### Sex-biased gene expression and population differentiation

Sayadi et al. (2019) showed that the hallmarks of balancing selection, presumably through sexually antagonistic pleiotropy, is particularly pronounced in genes with intermediate female bias in expression in *C. maculatus*. This was true for both within-population indices of selection and the degree of shared polymorphism across populations. They also demonstrated that strong male and female bias in expression are both associated with signs of weak selection, presumably because the efficacy of directional selection is reduced in genes with sex limited or sex biased expression (Dapper and Wade 2020). We therefore also assessed the pattern of population differentiation across sex biased gene expression, for the subset of genes where data on sex specific expression is available (Immonen et al. 2017).Although relative differentiation did tend to be higher for genes with female bias in expression, this was not significant for either population comparison (Figure 5).

**Figure 5.**
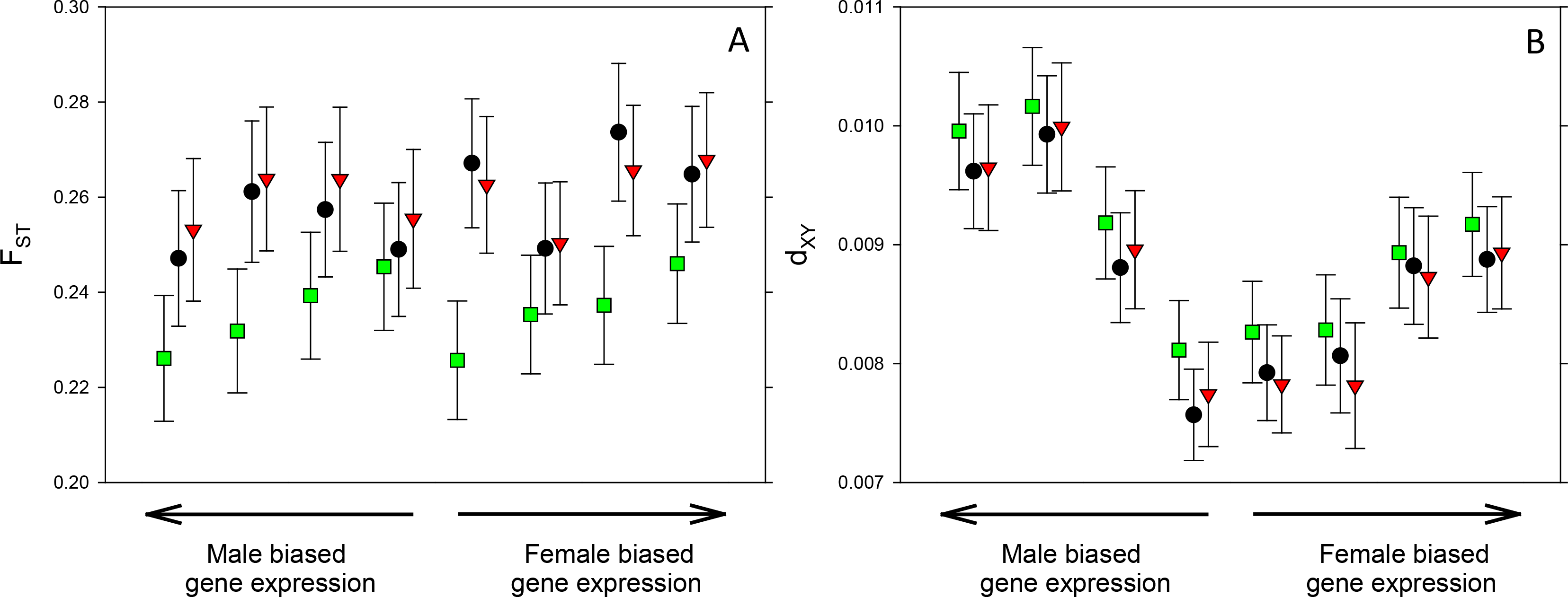
The role of sex biased gene expression for population divergence in three *C. maculatus* populations. Shown are mean (±95% bootstrap CI) (A) relative and (B) absolute population differentiation for pairs of populations for genes showing different degrees of sex-biased expression (green: C vs. Y; black: B vs. C; red: B vs. Y). Genes were grouped into quartiles based on their log2FC value, separately for FBGs and MBGs, resulting in eight bins in total. Sample size per bin is n = 591–656 genes. (A) Relative divergence tended to be highest for female biased genes and intermediately male biased genes, but the effect of gene expression was not significant in either population comparison (logit F_ST_; C vs. Y: F_7,4956_ = 2.02, *P* = 0.053; B vs. C: F_7,4956_ = 1.47, *P* = 0.177; B vs. Y: F_7,4956_ = 1.57, *P* = 0.146). (B) In constrast, both female biased and especially male biased genes showed considerably and significantly higher absolute divergence relative to genes with less sex biased expression (log10 dXY; C vs. Y: F_7,4985_ = 10.67, *P* < 0.001; B vs. C: F_7,4985_ = 10.50, *P* < 0.001; B vs. Y: F_7,4985_ = 9.74, *P* < 0.001) (*P* values from permutation tests).

Genes with sex biased expression, especially male biased expression, did show considerably and significantly higher absolute divergence (Figure 5). This could be interpreted as being consistent with a dominant role for drift in absolute differentiation between the populations studied here (Dapper and Wade 2020). However, we note that when adding population specific *π*N/*π*S ratios as covariates to models of the effect of sex bias in gene expression on d_XY_ (Figure 5), the effects of sex bias in gene expression remained highly significant (P < 0.001 in all three cases). This implies that properties of sex biased genes other than a lower efficacy of selection contributes to the higher level of absolute differentiation seen in sex biased genes. One possible interpretation is that female biased genes to larger extent show shared polymorphism and lower absolute differentiation due to balancing selection, while male biased genes to a larger extent contribute to absolute population differentiation through divergent positive selection. In line with this interpretation, Sayadi et al. (2019) found that female biased genes tend to show overall positive but male biased genes negative Tajima’s D within these populations.

### Effect size and fixation

Testing prediction VIII is greatly complicated by the fact that we lack explicit measures of the phenotypic effect size of segregating variants, as well as the fact that we lack detailed information of ancestral allele frequencies. To enable at least a provisional test of this prediction, we (i) assume that the number of populations in which a given gene shows non-synonymous polymorphism is inversely related to its fixation probability and (ii) lean on the common but perhaps somewhat dubious assumption that the level of gene expression is overall positively associated with size of allelic effects in that locus across protein coding genes (e.g. Duret and Mouchiroud 2000; Gout et al. 2010). We then compared the relationship between gene expression (i.e., overall FPKM values from Immonen et al. 2017) and within-population fixation for non-synonymous sites in genes involved in regulating metabolic processes (Biological Process GO:0019222) with that in all other genes (see Figure 6). This comparison is relevant given the prediction that metabolic rate should be a polygenic trait under stabilizing selection in our system. We found that both expression level (LLR_8_ = 15.35, P < 0.001) and gene class (LLR_1_ = 16.45, P < 0.001) had significant effects on fixation (Generalized Linear Model [GLIM], with Poisson errors, a log link and an empirical dispersion factor). More importantly, however, the interaction between expression level and gene class was significant (LLR_8_ = 3.13, P = 0.002). Genes involved in the regulation of metabolic processes tended to show the highest probability of fixation for intermediately expressed genes while other genes showed the highest probability of fixation for highly expressed genes (Figure 6), which is at least consistent with prediction VIII. A more complex GLIM, using mean centered log FPKM and its squared term as continuous predictors along with gene class, confirmed that the curvature of the pattern visualized in Figure 6 differed significantly for the two gene classes (gene class × FPKM^2^; LLR_1_ = 3.87, P = 0.049) and was convex for metabolic rate genes but concave for other genes.

**Figure 6.**
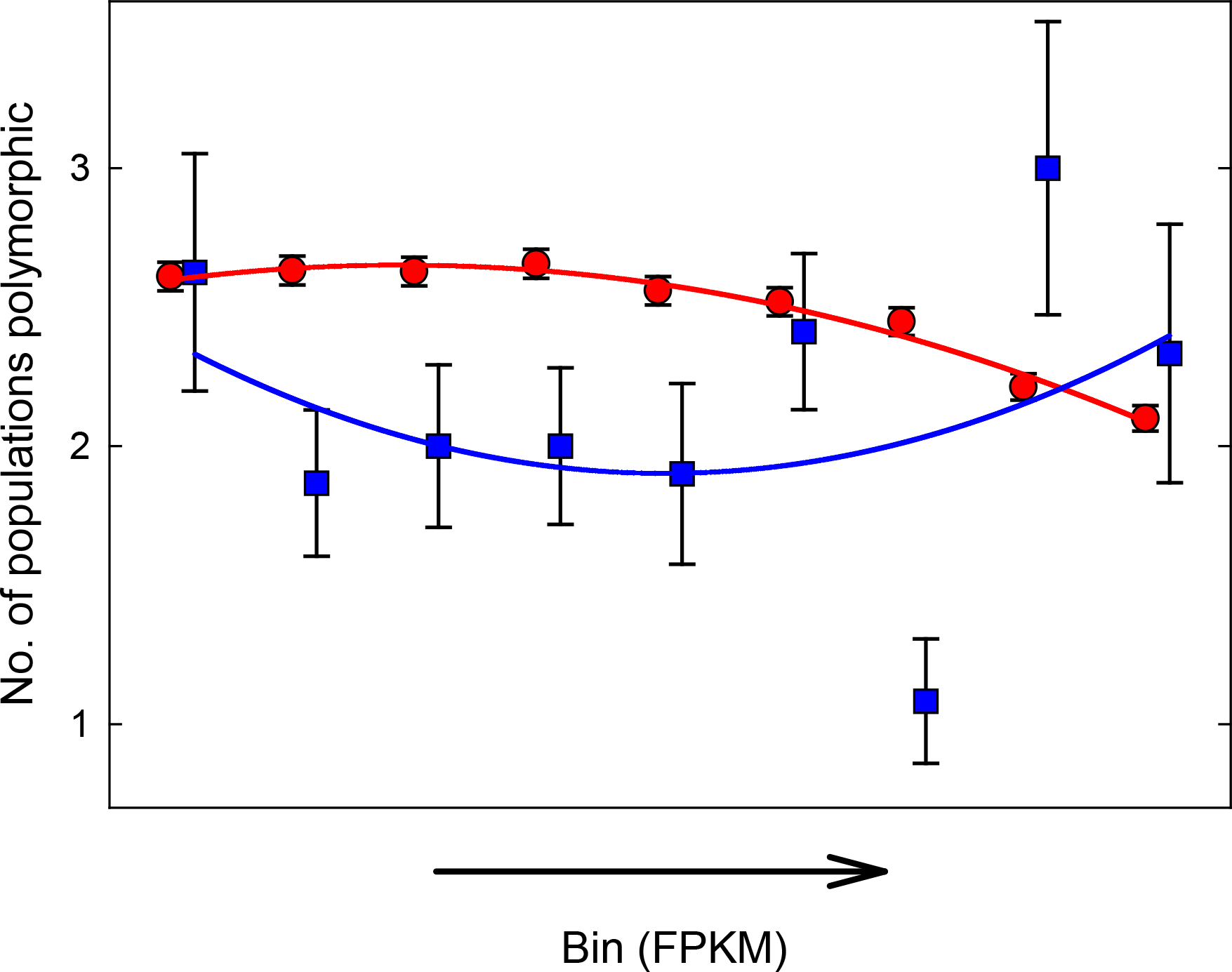
The number of populations in which genes show non-synonymous within-population SNPs as a function of increased expression level. Shown are predicted mean (±SE) polymorphism for (A) genes involved in the regulation of metabolic processes (blue squares) and (B) all other genes (red circles). Genes were grouped into nine equally sized bins (N = 552-556 genes per bin), based on their FPKM value. Note that the probability of fixation is assumed to be inversely related to the ordinate, such that more highly expressed non-metabolic genes were more likely to be fixed for non-synonymous sites within populations. Genes involved in the regulation of metabolic processes showed a significantly different pattern, with highest degree of fixation for intermediately expressed genes. Curves represent fitted squared functions (see text).

### Linked selection

Although the overall pattern described by the above analyses is robust, more detailed inferences about the significance of various terms potentially suffer from the common implicit, but unrealistic, assumption that genes segregate independently during divergence. However, the facts that (1) *C. maculatus* has a fairly large (1.2 gigabases) and repeat-rich (70%) genome with ten chromosome pairs (2n = 18 + XX/XY), (2) the population divergence is fairly deep (>3000 generations) and differentiation substantial (F_ST_’s 0.25) and (3) GC content, often used as a proxy for recombination rate (e.g. Rousselle et al. 2019), was not a strong predictor of gene specific differentiation, all suggest that linked selection is likely of relatively minor importance in our genome wide analyses. However, we lack a detailed recombination map of the *C. maculatus* genome and it is therefore not possible to explicitly model if and how variation in recombination rate across the genome affects divergence. The fact that the genome assembly, although showing a high fraction of well-assembled genes, is highly fragmented (Sayadi et al. 2019) also precludes scans of genomic islands of differentiation (Cruickshank and Hahn 2014; Burri 2017).

## Discussion

Our main aim was to apply an integrative and phenotypically informed “magnifying glass” approach (sensu Via 2009) to population genomic data in an attempt to shed some light on the role of stabilizing selection on polygenic life history traits in the evolution of incipient speciation in an insect. Although this is clearly a very challenging task, the patterns documented were well aligned with the predictions made. Overall, thus, we suggest that our findings are consistent with a scenario whereby populations have diverged in polygenic life history traits under stabilizing selection, which we suspect is the likely cause of the emerging reproductive isolation seen between the populations studied here (Figure 1). Yet, we note that a large number of interacting factors and processes should affect population divergence and it currently not possible to firmly assess how unique our predictions are for the scenario considered. Our approach and analyses did generate a series of insight. To us, two main findings stood out.

First, imbalances in inferred selection (i.e., positive Tajima’s D in one population and negative in the other) were positively related to F_ST_ across genes. As detailed in the introduction, we suggest that this pattern is predicted for divergence in polygenic traits under stabilizing selection. It is, however, clearly very difficult to entirely exclude the possibility that other processes and effects might also be capable of generating, or at the very least contributing to, this empirical pattern and it is currently not possible to fully evaluate this prediction. For example, there may be intrinsic mathematical dependencies between F_ST_ and properties of the underlying allele-frequency distributions (Jakobsson et al. 2013) that could contribute to complex and non-intuitive relationships between the two. Dedicated modelling efforts would be required to fully evaluate this possibility. Such simulation would, however, be complex and would need to explore variation in a large number of interacting factors and processes (see below) and are beyond the scope of our current contribution. Similarly, genetic drift is predicted to generate higher F_ST_ for loci with higher initial minor allele frequencies, as the Wright-Fisher diffusion process is contingent upon the binomial variance. Loci with more balanced polymorphism at the time of divergence may thus show higher F_ST_ after drift and may simultaneously tend to show an excess of rare variants in one population but retain a more even distributions of alleles in the other. These sorts of effects are, again, very difficult to account for, but we note that (i) entering *π*_N_/*π*_S_ to our models to account for the strength of positive selection did not reduce the effects of the interaction between Tajima’s D in our models, (ii) *π*_N_/*π*_S_ was negatively, rather than positively, related to F_ST_ and (iii) there was a striking and significant functional enrichment of genes showing high F_ST_ and an imbalance in Tajima’s D. These observations are, collectively, difficult to reconcile with a dominating role for random genetic drift in generating the pattern seen.

In addition, we expect the relationship between within-population site frequency spectra and between-population polygenic differentiation to depend on a large number of factors, such as, for example, time since divergence, mutation rate, recombination rate, effective population size, demography, number of genes involved, distribution of allelic effects, strength of phenotypic selection and differences in optima (Barton 1986, 1989; Fierst and Hansen 2010; Höllinger et al. 2019; Barghi et al. 2020). There might thus be conditions where the predicted pattern is most apparent, but it is unclear exactly what those are and explicit modelling efforts aimed at clarifying this would be challenging but helpful. It is also not clear whether the genome wide pattern that we document is commonly observed among other potential empirical examples of polygenic population divergence. Although a number of studies of genome wide divergence have estimated both F_ST_ and Tajima’s D, few have explicitly related within-population Tajima’s D to between-population F_ST_ and among those that have (e.g. Franks et al. 2016) none seem to have modelled the interactive effect predicted and observed here.

Second, phenotype-free bottom-up approaches in speciation genomics suffer from well-known limitations, in particular when adaptive divergence involves polygenic traits under stabilizing selection (e.g. Rockman 2012; Korte and Farlow 2013; Wellenreuther and Hansson 2016; Hayward and Sella 2021). Several authors have suggested that vertical integration of information and data across different levels of biological organization can help alleviate some of these limitations (e.g. Lee at el. 2014; Seehausen et al. 2014; Pardo-Diaz et al. 2015). For example, phenotypically informed genomic approaches, where an understanding of phenotypic selection is used to generate a priori predictions against which genomic data is analyzed and interpreted, can enrich and improve our inferential basis (Pavlidis et al. 2012; Savolainen et al. 2013). For polygenic traits, Hayward and Sella (2021) suggested that we should perhaps aim to identify traits, rather than specific loci, showing adaptive divergence. Based on a large body of work on phenotypic selection and trait divergence within and between *C. maculatus* populations, we predicted that polygenic life history traits that are determined by metabolic processes should be at the center of population divergence and that such traits should be under net stabilizing selection. Our results match these predictions, as (i) genes showing high F_ST_ and an imbalance in Tajima’s Ds were enriched with genes involved in metabolic processes, (ii) this enrichment was repeatable over population comparisons and (iii) genes known to be involved in the regulation of metabolic rate showed both a higher F_ST_, a strong association between imbalance in Tajima’s Ds and F_ST_ and a distinct pattern of fixation as a function of gene expression. In this sense, our findings can be seen as illustrating that a thorough understanding of phenotypic selection can be used to inform inferences based on genomic data.

Needless to say, using a phenotypically informed approach might be difficult in many systems, where detailed information on phenotypic selection and divergence in life history traits is lacking. Yet, even in systems where little a prior information on phenotypic selection and divergence is available, some general predictions could be made. For example, one would expect that metabolic genes should often stand out. This is because metabolic networks are inherently polygenic and because we generally expect net selection on metabolic fluxes to be stabilizing (Whitlock et al 1995). Similar predictions could perhaps be made about genes involved in other complex processing, developmental or signaling pathways (Csilléry et al. 2018).

Gene specific absolute and relative differentiation quantify different aspects of differentiation (Cruickshank and Hahn 2014) and several studies have found that they are at most only weakly related (e.g. Irwin et al. 2016; van Doren et al. 2017; Vijay et al. 2017; Talla et al. 2019). Our findings parallel these studies. Moreover, we found that d_XY_ showed a distinct signal in terms of how it related to inferred selection within populations. We found that d_XY_ was consistently highest for genes showing signs of less constraint (higher *π*N/*π*S) and allele frequency spectra more consistent with neutrality (near zero Tajima’s D_NS_). In addition, genes with a generally high d_XY_ across all population comparisons (i.e., high PC1_dXY_) were those with high *π*N/*π*S and this set showed no apparent enrichment in terms of gene function, which also suggests that stochastic processes may be major contributors to absolute differentiation. Interestingly, the relatively high repeatability of d_XY_ across the three population comparisons would then imply a lower, rather than higher, redundancy for genes showing less constraint. On the one hand, thus, our collective findings confirm an important role for drift in absolute population differentiation. On the other hand, they also suggest that those genes that show high absolute differentiation may not generally be the most important contributors to incipient genetic incompatibility and reproductive isolation, by the mere fact that they tend to show signs of neutrality.

We also compared differentiation across certain focal functional classes of genes, using annotations based on experimental data in *C. maculatus* (Bayram et al. 2017; Bayram et al. 2019; Sayadi et al. 2019). We were particularly interested in estimating the relative potential roles of sexual selection and natural selection in driving population divergence. Sexual selection in *C. maculatus* is to a large extent generated by postmating sexual selection (Fritzsche and Arnqvist 2013) and a key male phenotype in mediating competitive fertilization success among males is the ejaculate proteome (Yamane et al. 2015). Males transfer more than 300 proteins to females during mating (Bayram et al. 2019), many of which are accessory gland proteins, and compositional variation in this multidimensional phenotype is striking: the relative abundance of seminal fluid proteins varies markedly across populations and is also related to male fertilization success (Goenaga et al. 2015). Seminal fluid protein genes are thus candidate genes for sexual selection in males. Yet, Sayadi et al. (2019) found little evidence for selection on seminal fluid genes within populations, apart from a signal of stronger purifying selection, and our study shows that these genes generally only show a slightly elevated rate of relative differentiation between populations. They did, however, consistently show a high absolute differentiation (SI Table 3).This may at least partly reflect functional differences rather than drift and thus contribute to reproductive isolation, given that seminal fluid proteins are known to be functionally important for male reproductive success in this species (Goenaga et al. 2015; Yamane et al. 2015). In agreement with this, and unlike in the entire gene set, PC1_dXY_ did not show a positive relationship with *π*N/*π*S in the three populations across all seminal fluid protein genes (multiple regression; *F*_3,100_ = 1.29, *P* = 0.282; two out of three *β'*s even negative) suggesting that drift does not generally promote divergence in male seminal fluid genes. Taken together, these findings are somewhat inconclusive. Male reproductive proteins are known to often evolve rapidly (e.g. Avila et al. 2011) and while it seems very likely that some seminal fluid proteins indeed diverge rapidly as a result of divergent positive sexual selection in *C. maculatus*, this is seemingly manifested as absolute rather than relative differentiation at the scale investigated here.

Genes encoding enzymes that larvae employ to detoxify and digest legume host seeds are prime candidate genes for host adaptation and natural selection in these herbivorous insects (Zhu-Salzman and Zeng 2015). We found that genes encoding enzymes with digestive function in the larval gut showed markedly elevated relative, but not absolute, differentiation compared to other genes (SI Table 3). This is in line with a key role for natural selection in population divergence. We conclude that our findings are consistent with important roles for both sexual and natural selection in divergence, but our results also suggest that the mode of selection and differentiation may differ for the two classes of candidate genes inspected here. It is currently unclear, however, whether and how such differences translate into differences in their importance in terms of generating reproductive isolation.

This study represents an attempt to evaluate the role of selection for polygenic population divergence that extends conventional assessments of the role of selective sweeps involving independently acting genes. Although some of the patterns documented here can clearly have alternative or complimentary explanations, we feel that our collective findings are encouraging and suggest that polygenic adaptation may leave a detectable footprint in the covariance between allele frequency spectra within populations and divergence between populations. We found that relative population differentiation was to a large extent built by genes showing excess of rare variants in one population and simultaneously a more even distribution of allele frequencies in the other. Importantly, these genes were enriched with genes with a metabolic function and this enrichment was repeatable across population comparisons. Across many interacting loci, differences in allele frequencies are likely to result in BDM incompatibilities in crosses between populations, manifested as suboptimal metabolic phenotypes. We suggest that more genome wide studies that explicitly relate the properties of within-population allele frequency spectra to between-population divergence against a phenotypically informed backdrop are worthwhile, in that they may provide insights into the genetics of incipient speciation. Yet, it is also clear that more efforts to explicitly model these scenarios are needed to more firmly identify the empirical patterns expected.

## Materials and Methods

### Populations

The populations of the seed beetle *C. maculatus* used here were originally collected in three different geographic regions; California (1980), Yemen (1976) and Brazil (2000). They were all founded by a large number of individuals (>>100) and have since been maintained in the laboratory on their natural host *V. unguiculata* at population sizes of >500 individuals and at 29°C, 60-70% RH and a 12L:12D light cycle. Demographic modelling of genetic variation in natural populations have shown that effective population size is very large in natural populations *C. maculatus* (Kebe et al. 2017), consistent with the fact that it is a cosmopolitan and very abundant crop pest, but inbreeding in laboratory populations is always a potential concern. Four lines of evidence suggest that this is likely not the case in these particular populations. First, quantitative genetic studies suggests that laboratory populations of seed beetles such as these do not suffer from inbreeding depression but harbor levels of genetic variation typical of that of natural populations (Fox et al. 2007). Second, molecular genetic estimates of effective population size in similar laboratory seed beetle populations suggest that effective population size is fairly large (N_e_ > 1000 individuals; Gompert and Messina 2016). Third, and perhaps most importantly, the mean within-population genome-wide SNP density in coding regions across all genes is approximately 6 SNPs / kb in the studied populations and the level of nucleotide diversity for synonymous sites in all three populations is *π*_S_ ≈ 0.022 (Sayadi et al. 2019). This is comparable to, or even larger than, that found in natural population of seed beetles (Tine et al. 2013), as well as in many other insects (e.g. Gossman et al. 2012; Chen et al. 2017), and is inconsistent with significant effects of inbreeding (Charlesworth 2003).

In studies of divergence between laboratory populations, admixture in the laboratory is a common concern for species like *Drosophila melanogaster* where adults are strong fliers. However, adult *C. maculatus* are poor fliers and very rarely take to the wings in the laboratory and larvae are contained to a single host bean throughout their development. These factors preclude significant admixture, which is consistent with a large number of private SNPs observed in these populations (Sayadi et al. 2019) and the high degree of differentiation seen. Further, effects of recent admixture on the signal of differentiation would tend to be localized to certain regions of the genome, which we did not observe here (see below).

*Callosobruchus maculatus* was brought to Asia from West Africa by human farmers of its main host about 2,000 years ago and were introduced to the Americas from West Africa by Spanish settlers in the early 1700s (Kebe et al. 2017). This suggests that Yemen originally diverged from ancestral African populations some 20,000 generations ago while California and Yemen diverged some 3,000 generations ago. Because the climate differs substantially in the three different regions, the source populations are expected to show local adaptation to differences in temperature and humidity and theory suggest that these divergence times are likely deep enough for the populations to have achieved the peak shifts and have reached an equilibrium phase (Hayward and Sella 2021). However, the degree of adaptive genetic differentiation has most likely been restricted by gene flow resulting from commercial trade with its host (Kebe et al. 2017). Genetic differentiation between these laboratory populations should thus certainly to some extent reflects differences in life history optima in their parental populations, but also the effects of gene flow, genetic drift, quasi-neutral polygenic drift and adaptation to common garden conditions in the laboratory. While our data thus does not allow us to estimate the precise extent to which divergence has evolved in the wild and in the laboratory, it is still useful for testing the utility of our approach for detecting a role of peak shifts in polygenic population divergence.

### Sequencing and analyses

All sequencing data derives from Sayadi et al. (2019) and we refer in full to that study for details on read data and data availability. Briefly, to assess variation within and divergence between populations, we sequenced pools of individuals from the three populations (n = 200 males per population), to an average depth of ~125X per population using the Illumina HiSeq2500 system with a paired-end 125bp read length. Here, we focus on gene and population specific estimates of Tajima’s D for non-synonymous sites (SI Figure 4) and for *π*N/*π*S which we extracted using popoolation and popoolation2 (Kofler et al. 2011a, 2011b). Tajima’s D was standardized and *π*N/*π*S ratios log10 transformed prior to statistical modelling. Gene specific relative divergence (F_ST_) was estimated using the unbiased method proposed by Hivert et al. (2018) as implemented in poolfstat, using default options. Absolute divergence (d_XY_) was estimated as A_X_B_Y_ + A_Y_B_X_, where A and B are the frequencies of two variants at a given site and X and Y represent the two populations being compared (Delmore et al. 2015; Dennenmoser et al. 2017), and gene specific estimates were derived by summing over the gene (i.e., the window) and dividing by the total number of sites (variable or not) (see Van Doren et al. 2017). To normalize residuals, F_ST_ was logit transformed and d_XY_ log10 transformed prior to statistical modelling.

To identify overrepresentation of Gene Ontology terms, we used a hypergeometric test with a *P*-value cutoff < 0.05 as implemented in the GOstats package v.2.46.0 (Falcon and Gentleman 2007). Gene universes are defined in SI Table 1.

### Gene classes

Because certain classes of genes are of particular interest in terms of population divergence and incipient speciation, we also explored differentiation in a few focal classes of genes. These were genes encoding (i) male ejaculate proteins (Bayram et al. 2017, 2019), (ii) female reproductive proteins (Bayram et al. 2019), (iii) enzymes involved in digestion and food processing in larvae and (iv) genes located on the sex chromosomes (Sayadi et al. 2019).

## Supplemental Information

Supplementary figures and tables are available at *XXXX*.

## Acknowledgements

This work was supported by grants the European Research council ERC (GENCON AdG- 294333 to GA), the Swedish Research Council VR (621-2014-4523 to GA) and FORMAS (2018- 00705 to GA). We are grateful to N. Barton, T. Hansen and the GENCON lab group for constructive discussions and to E. Arnqvist for support. The computations and data handling were enabled by resources provided by the Swedish National Infrastructure for Computing (SNIC) at UPPMAX partially funded by the Swedish Research Council, through grant agreement no. 2018-05973.

## Author Contributions

G.A. conceptualized the study and acquired funding. A.S. performed all bioinformatic analyses, while G.A. performed statistical modelling. G.A. wrote the original draft. Both authors edited the manuscript.

## Data Accessibility

The annotated genome assembly of *C. maculatus* used here (Sayadi et al. 2019) is available from the European Nucleotide Archive under accession PRJEB30475. Pool-seq raw sequencing data have been deposited at the NCBI sequence read archive, under the accession number PRJNA503561.

