## Supplemental Information for "A possible genomic footprint of polygenic adaptation on population divergence in seed beetles?"

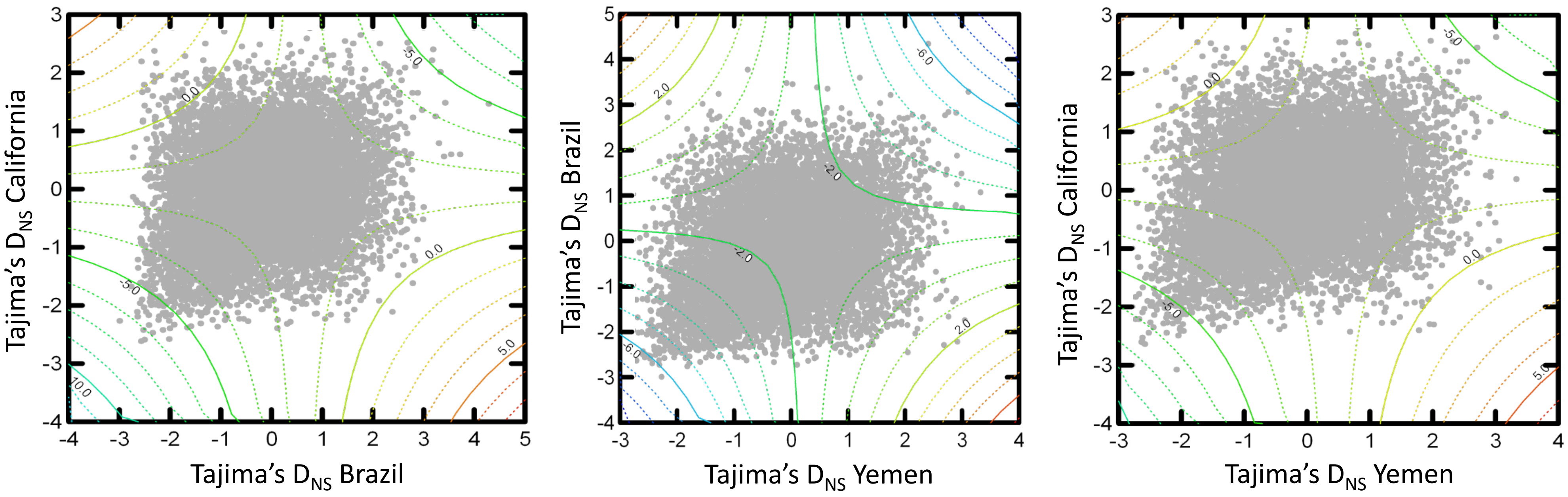


SI Figure 1. Scatterplots of Tajima's D (based on non-synonymous SNPs) between pairs of populations across all genes (r = 0.23 – 0.27, $P < 10^{-14}$ in all three cases). Contours represent predicted $F_{ST}$ on a logit scale from linear models including Tajima's D in the two populations, their quadratic term and their interaction (i.e., model A1, B1 and C1 in Table X). On this scale 0.0 corresponds to $F_{ST}$ = 0.5, 1.0 [-1.0] corresponds to $F_{ST}$ = 0.91 [0.09] and 2.0 [-2.0] corresponds to $F_{ST}$ = 0.99 [0.01]. The 3D saddle surface in the *Z*-dimension illustrates the fact that genes showing a signal of balancing selection in one population (i.e., positive Tajima's D) and positive in the other (i.e., negative Tajima's D) show the highest overall $F_{ST}$ (upper left and lower right hand corners). The comparison B – C is shown to the left, B – Y in the centre, and C – Y to the right.

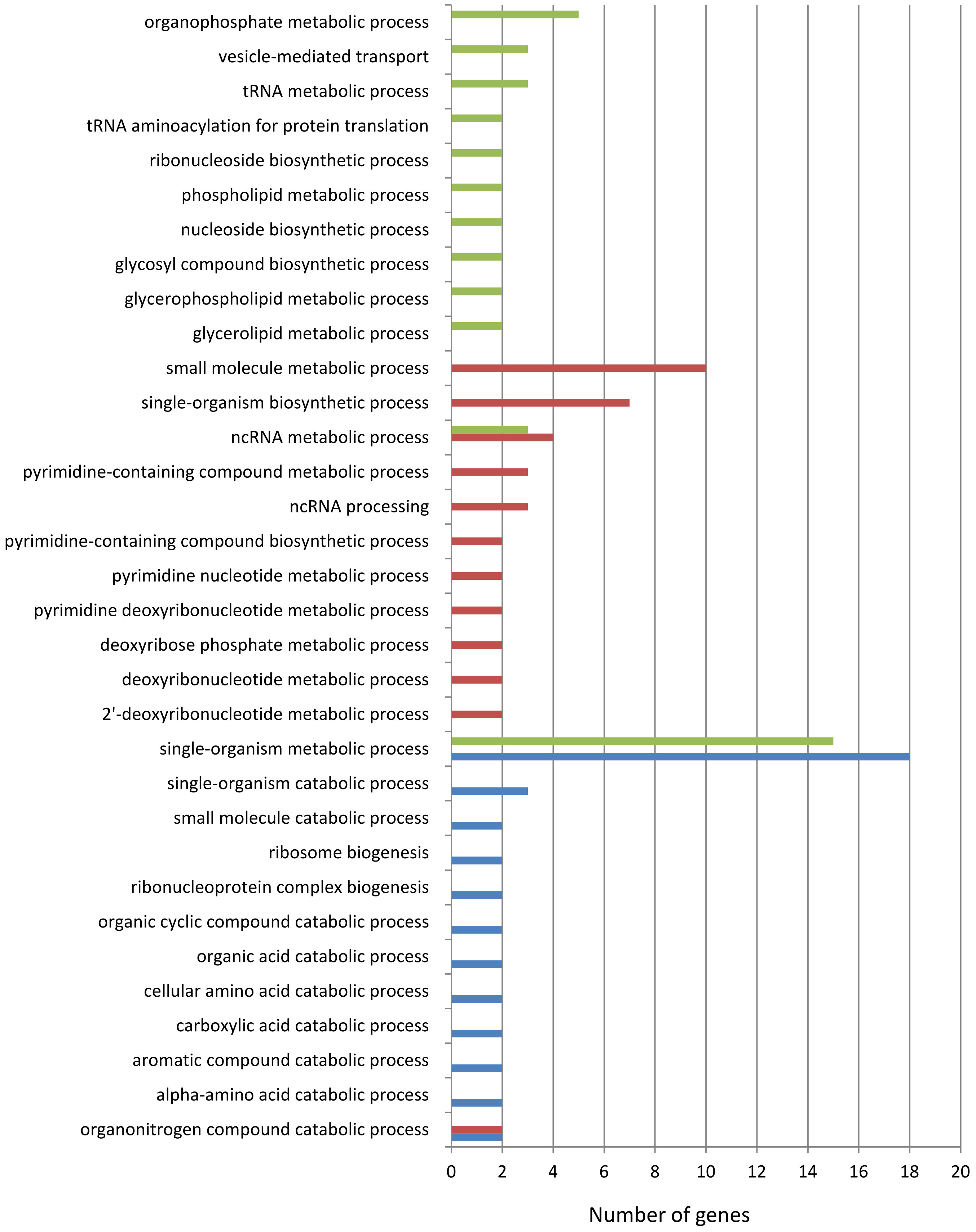


SI Figure 2. Gene ontology (GO) term enrichment analysis (BP) of genes showing an imbalance in Tajimas $D_{NS}$ and a $F_{ST} > 0.3$. All terms shown are significant (at $P < 0.05$) and contain at least 2 genes (green: C vs. Y; red: B vs. Y; blue: B vs. C). Note the consistent enrichment of metabolic, catabolic and biosynthetic processes.

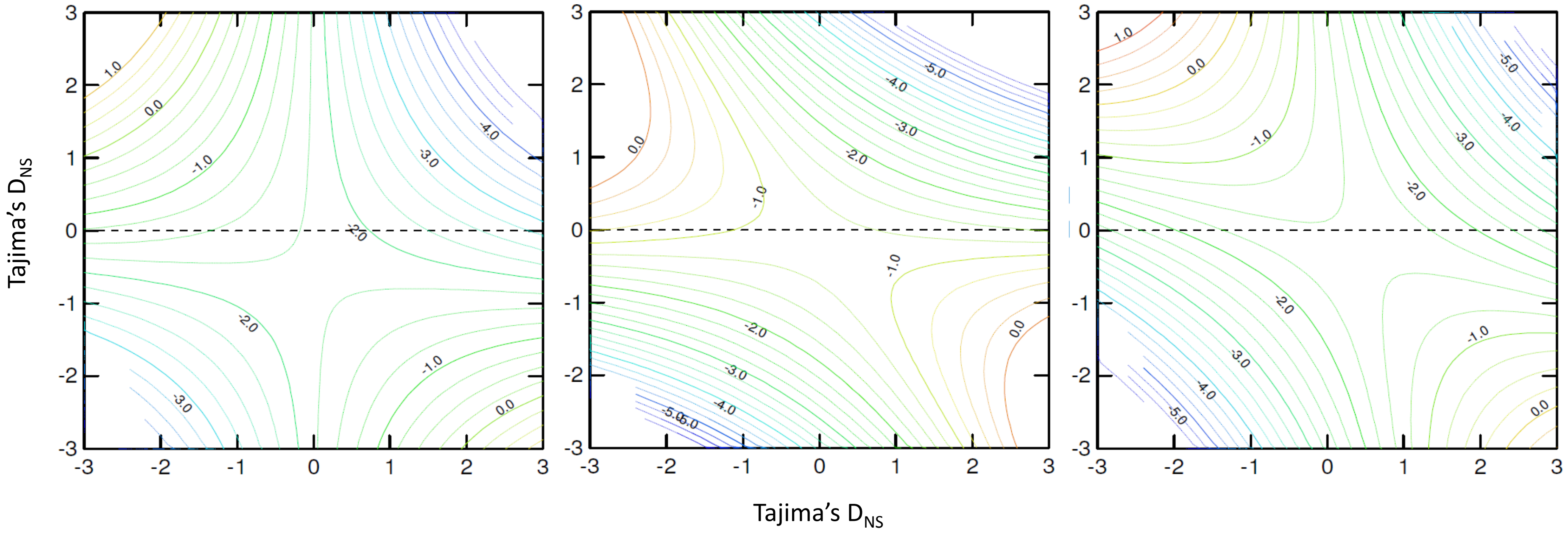


SI Figure 3. Contour plots from full response surface models (Myers et al. 2016) of $D_{NS}$ in two populations on predicted logit $F_{ST}$ (*Z* dimension) between them, including only genes annotated as being involved in processes that regulates the rate of metabolic organismal pathways (N = 116) (left: B vs. C; center: B vs. Y; right: C vs. Y). As in models involving all genes, interaction effects were very strong (SI Table X) and the the fitted function describes a marked saddle surface. Again, genes with highest predicted $F_{ST}$ were those with negative Tajima's D in one population and positive in the other. Logit $F_{ST}$ = 0 corresponds to $F_{ST}$ = 0.5.

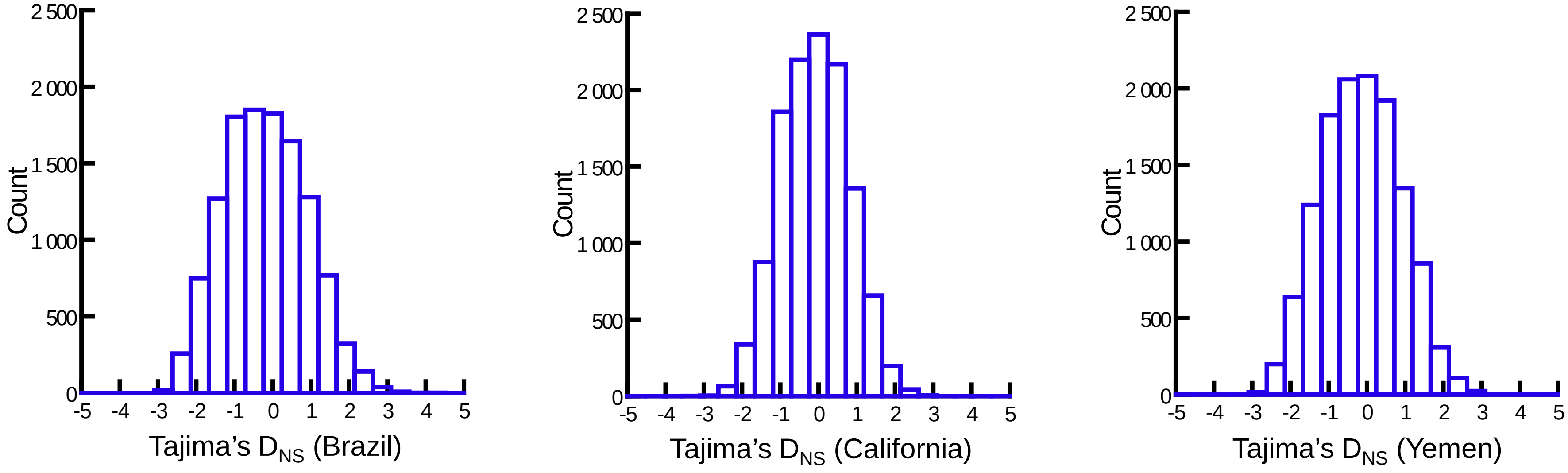


SI Figure 4. Frequency distribution of Tajima's D for non-synonymous sites across all genes in the three populations of *C. maculatus* studied. Mean (SD) Tajima's $D_{NS}$ was B: -0.192 (1.07), C: -0.093 (0.87), Y: -0.150 (1.01).

SI Table 2. Analyses of variance of the full surface response models of the effects of Tajima's D (standardized) within two populations on logit $F_{ST}$ between them, across all genes annotated as being involved in regulation of metabolic rate. The effect of the interaction between Tajima's D in the two populations on $F_{ST}$ across these sets of genes was very strong relative to the linear and quadratic effects (see F-ratios). See SI Figure 3 for visualizations of the full surface response models.

| | **Source** | **df** | **SS (Type I)** | **F-ratio** | **P** |
|---|---|---|---|---|---|
| B vs. Y | Linear | 2 | 3.58 | 0.82 | 0.445 |
| | Quadratic | 2 | 22.12 | 5.05 | 0.008 |
| | Interaction | 1 | 22.63 | 10.32 | 0.002 |
| | Residual | 106 | 232.35 | | |
| | Total | 111 | 280.67 | | |
| B vs. C | Linear | 2 | 9.51 | 1.42 | 0.247 |
| | Quadratic | 2 | 3.79 | 0.57 | 0.570 |
| | Interaction | 1 | 12.41 | 3.70 | 0.057 |
| | Residual | 105 | 351.92 | | |
| | Total | 110 | 377.63 | | |
| C vs. Y | Linear | 2 | 3.30 | 0.72 | 0.489 |
| | Quadratic | 2 | 16.02 | 3.50 | 0.034 |
| | Interaction | 1 | 22.49 | 9.82 | 0.002 |
| | Residual | 110 | 251.78 | | |
| | Total | 115 | 293.58 | | |

**SI Table 3.** The results of general linear models of differences in mean differentiation constrasting sets of genes with the remainder of all genes. Analyses were performed on logit transformed $F_{ST}$ and log10 transformed $d_{XY}$ and *P*- values derive from permutation tests (9999 randomizations).

| | $F_{ST}$ | | | | | | $d_{XY}$ | | | | | |
|---|---|---|---|---|---|---|---|---|---|---|---|---|
| | B vs. C | | B vs. Y | | C vs. Y | | B vs. C | | B vs. Y | | C vs. Y | |
| Gene set | $F_{ndf,ddf}$ | *P* | $F_{ndf,ddf}$ | *P* | $F_{ndf,ddf}$ | *P* | $F_{ndf,ddf}$ | *P* | $F_{ndf,ddf}$ | *P* | $F_{ndf,ddf}$ | *P* |
| Male seminal fluid proteins (N = 184) | **$4.37_{1,15306}$** | **0.048** | $3.92_{1,15306}$ | 0.083 | $3.90_{1,15306}$ | 0.090 | **$9.35_{1,19064}$** | **0.005** | **$4.73_{1,19178}$** | **0.045** | **$9.16_{1,19097}$** | **0.003** |
| Female reproductive proteins (N = 126) | $0.47_{1,15305}$ | 0.547 | $3.45_{1,15305}$ | 0.104 | $0.15_{1,15305}$ | 0.725 | $0.26_{1,19063}$ | 0.612 | $0.01_{1,19177}$ | 0.930 | $0.01_{1,19096}$ | 0.934 |
| Digestive enzymes (N = 740) | **$13.99_{1,15305}$** | **<0.001** | **$20.62_{1,15305}$** | **<0.001** | **$12.93_{1,15305}$** | **0.002** | $2.28_{1,19063}$ | 0.151 | $3.68_{1,19177}$ | 0.066 | $3.53_{1,19096}$ | 0.075 |
| Candidate X-linked genes (N = 604) | **$19.32_{1,15305}$** | **<0.001** | **$28.55_{1,15305}$** | **<0.001** | **$12.77_{1,15305}$** | **0.003** | **$385.96_{1,19063}$** | **<0.001** | **$484.21_{1,19177}$** | **<0.001** | **$378.94_{1,19096}$** | **<0.001** |
| Candidate Y-linked genes (N = 277) | **$8.23_{1,15305}$** | **0.015** | **$47.79_{1,15305}$** | **<0.001** | **$22.54_{1,15305}$** | **<0.001** | **$781.73_{1,19063}$** | **<0.001** | **$1002.03_{1,19176}$** | **<0.001** | **$831.90_{1,19096}$** | **<0.001** |
